# Characterization of uranyl (UO_2_^2+^) ion binding to amyloid beta (Aβ) peptides: effects on Aβ structure and aggregation

**DOI:** 10.1101/2023.03.29.534802

**Authors:** Elina Berntsson, Faraz Vosough, Andra Noormägi, Kärt Padari, Fanny Asplund, Maciej Gielnik, Suman Paul, Jüri Jarvet, Vello Tõugu, Per M. Roos, Maciej Kozak, Astrid Gräslund, Andreas Barth, Margus Pooga, Peep Palumaa, Sebastian K. T. S. Wärmländer

## Abstract

Uranium (U) is naturally present in ambient air, water, and soil, and depleted uranium (DU) is released into the environment via industrial and military activities. While the radiological damage from U is rather well understood, less is known about the chemical damage mechanisms, which dominate in DU. Heavy metal exposure is associated with numerous health conditions including Alzheimer’s disease (AD), the most prevalent age-related cause of dementia. The pathological hallmark of AD is deposition of amyloid plaques, consisting mainly of amyloid-β (Aβ) peptides aggregated into amyloid fibrils in the brain. However, the toxic species in AD are likely oligomeric Aβ aggregates. Exposure to heavy metals such as Cd, Hg, Mn, and Pb is known to increase Aβ production, and these metals bind to Aβ peptides and modulate their aggregation. Possible effects of U in AD pathology have been sparsely studied. Here, we use biophysical techniques to study *in vitro* interactions between Aβ peptides and uranyl ions, UO_2_^2+^, of DU. We show for the first time that uranyl ions bind to Aβ peptides with affinities in the micromolar range, induce structural changes in Aβ monomers and oligomers, and inhibit Aβ fibrillization. General toxic mechanisms of uranyl ions could be modulation of protein folding, misfolding, and aggregation.

## 1. Introduction

Alzheimer’s disease (AD) is the most common neurodegenerative disease among elderly people ^1-2^. The main AD risk factors are old age and genetic factors such as unfavourable alleles of the ApoE gene and sometimes Down’s syndrome ^1-2^, but also environmental risk factors such as smoking, diabetes, and traumatic brain injury ^2-4^. Identifying molecular targets to diagnose and treat the disease is imperative ^5-6^, and a number of drug candidates have been proposed with varying success ^7-10^.

The main molecular events underlying AD pathology appear to be aggregation of amyloid-β (Aβ) peptides and tau proteins into toxic soluble oligomers ^11-12^, and then into insoluble amyloid fibrils or tangles that deposit as plaques in the brains of AD patients ^2^, ^13^. The Aβ peptides are cleaved by β- and γ-secretase enzymes from the Amyloid-β precursor protein (APP) into peptides of different lengths ^14^, where Aβ(1- 40), i.e. Aβ_40_, and Aβ(1-42), i.e. Aβ_42_, are the most common ^15^. Both variants are unstructured monomers in aqueous solution but can adopt β-sheet ^16^ or α-helix ^17^ secondary structures in other environments ^18^. The β-sheet structure is of particular interest, as Aβ peptides in β-sheet hairpin conformations appear to be the building blocks of the aggregated fibrils ^16^.

The accumulation and aggregation of Aβ peptides can be influenced by a number of interacting factors ^3^, ^19^, such as other proteins and peptides ^10^, ^20-21^, including other amyloidic peptides/proteins ^22-25^, cationic molecules and metal ions ^26-28^, and various small molecules ^8^. Redox-active metal ions such as Cu(II) and Fe(II) are of special interest ^27^, ^29^, as they are present in the plaques of AD brains ^30-31^. For example, they can generate harmful oxygen radicals (reactive oxygen species; ROS) that may contribute to the AD pathology ^32-33^. Heavy metals are generally known to be toxic, but their toxic mechanisms are not fully understood ^34^. Known molecular mechanisms for metal toxicity include molecular and ionic mimicry ^35^, but other mechanisms are also possible, such as modulation of protein misfolding or aggregation ^36-38^. Thus, lead (Pb) ions not only compete with essential metals such as zinc and calcium ^39^, but they also increase the expression of APP and β-secretase ^40^, and Pb(IV) ions bind to Aβ peptides and modulate their aggregation ^3^. Similar effects on Aβ production and aggregation have been observed for the heavy metals mercury and cadmium ^3^, ^29^, ^40-43^.

Uranium (U) is a heavy metal with known neurotoxic effects ^34^, but these effects are sparsely studied, and a relation between U exposure and AD has not been established ^44^. However, the blood-brain-barrier does not prevent transfer of U into the central nervous system ^45^, and U has been shown to accumulate in the brain ^46^. Uranium has no biological function in the human body ^47-48^, and the adverse health effects of U exposure involve combined chemical and radiological mechanisms ^49-50^. Although the chemical toxicity is more severe ^51^, the radiological damage mechanisms are currently better understood ^34^, ^50^. The chemical toxicity obviously dominates in depleted uranium (DU), where the amount of the ^235^U isotope has been significantly reduced in favour of the less radioactive (i.e., longer half-life) ^238^U isotope ^47^. As DU is used in military equipment including certain ammunitions ^52^, DU contamination has emerged as a potential environmental problem in regions of war such as Iraq and the Balkans ^53-59^, with unclear health consequences for soldiers and civilians ^47^, ^49^, ^60-62^. To what extent DU contributes to leukemia or lung cancer remains debated ^63-66^.

Here, we use biophysical spectroscopic and imaging techniques to investigate the binding interactions between uranyl ions and the three Aβ peptide variants Aβ_40_, Aβ_42_, and the Aβ_40_(H6A, H13A, H14A) mutant. Of particular interest is the effect of uranyl on Aβ structure and aggregation.

## 2. Chemical, environmental, and toxicological aspects of uranium and uranyl ions

Uranium is present in ambient air, water, and soil. The highest human exposures result from drinking well water in geological regions rich in U ^34^, ^48^, ^67-70^. Regulatory Agencies have limited highest allowed concentration of U in drinking water to 30 µg/L ^71^, replacing a previous WHO limit of 15 µg/L. Absorption of U is low regardless of exposure route and highly dependent on its solubility ^34^. Inhaled U dust particles of low solubility can be retained in tissues for many years ^72^. Occupational exposure to U has historically involved miners handling uranium oxide, so called “yellowcake”, workers in the production of phosphate fertilizers and workers producing glazed pottery ^34^, ^73^. Sleep disturbances and possibly encephalitis has been linked to U exposure in former U mining districts in Kazakstan ^74^.

The main target for U toxicity is the kidney where atrophy and necrosis of glomerular walls have been noted ^34^, ^75-77^. Uranium also accumulates in bone ^34^, ^55^, ^78-80^. Uranium crosses the blood-brain-barrier ^45^ to accumulate in the central nervous system where inhalation and ingestion of U yields heterogeneous but specific accumulations, most prominent in the hippocampus region ^81^, responsible for memory recall. Rats surgically implanted with U pellets for 6 months have shown presence of U in the cortex, midbrain, and cerebellum ^82^. A study on mice showed toxic effects of depleted uranium in the mouse fetus ^83^, indicating that U can cross also the placental barrier.

There are currently no known correlations between uranium exposure and diseases such as leukemia ^68^, stomach cancer ^84^, liver or bladder cancer ^85^, or, as mentioned above, Alzheimer’s disease ^44^. One study however found cerebrospinal fluid U concentrations, albeit at low overall concentrations, to be significantly elevated in 17 patients with amyotrophic lateral sclerosis (ALS) when compared to 10 controls ^86^. Studies in rodents have related U exposure to distorted social behaviour ^87^ and weakened sensoriimotor behavior ^88^. Laboratory experiments using organisms such as rats and Caenorhabditis elegans have concluded with low acute neurotoxic potential of U following exposure, and a protective potential from the small metal-regulating protein metallothionein ^82^, ^89^. Protective effects have also been observed for the ghrelin hormone ^90^ and for antioxidant agents including glutathione ^91-92^

The most predominant, most stable, and most relevant form of uranium in aerobic environments is the oxycation uranyl, UO_2_^2+^. This ion is common in uranium-containing minerals, but it is also water-soluble and is present in the ocean at a concentration of 13.7 nM ^93^. The uranyl ion is paramagnetic ^94-95^, and in aqueous solution it acts as a weak acid with a pKa around 4.2 ^96^. It behaves as a hard acceptor and prefers to form complexes with fluoride and oxygen donor atoms, preferably in planar geometry involving four, five, or even six binding ligands ^97^. The capacity to accommodate five or six equatorial ligands in pentagonal or hexagonal bipyramidal coordination separates the uranyl ion from most other metal ions ^93^. Thus, these uncommon binding geometries have been employed in attempts to design uranyl-specific binding proteins, to e.g. extract uranium from sea water ^93^. As the most common form of uranium, most experimental studies of uranium toxicity have actually been conducted with uranyl ions rather than with metallic uranium, which is extremely rare in nature.

Interestingly, U intoxication and AD appear to have a common risk factor in the gene coding for the apolipoprotein E (ApoE). U exposure in mice was found to induce cognitive impairment in ApoE-deficient (ApoE−/−) males, together with some changes in cholesterol levels and metabolism ^98-99^. This is consistent with the ApoEε4 allele of the ApoE gene being a risk factor for mercury toxicity ^100-101^, and with ApoE-deficient mice showing increased iron accumulation in tissue over time ^102^. It therefore appears likely that the ApoE protein is involved in regulating metal homeostasis. The ApoEε4 allele is further linked to an increased probability of developing AD ^103-107^. Why the ApoE gene is a risk factor for both heavy metal toxicity and AD is currently unclear, and various explanations have been proposed ^103^, ^108-109^. The previously suggested hypothesis that ApoE might bind and transport metal ions via Cys residues ^100-101^, which are present in ApoE2 and ApoE3 but not in ApoE4, has recently been called into question ^110^.

## 3. Materials and Methods

### 3.1 Materials

Sodium dodecyl sulphate (SDS) detergent, 2-(N-morpholino) ethane sulfonic acid (MES) hydrate, sodium phosphate, depleted uranyl acetate, and dimethyl sulfoxide (DMSO) were all purchased from Sigma-Aldrich (USA). Synthetic lyophilized wildtype (wt) Aβ(1-42) peptides, abbreviated as Aβ_42_, with the primary sequence DAEFR_5_HDSGY_10_EVHHQ_15_KLVFF_20_AEDVG_25_SNKGA_30_IIGLM_35_VGGVV_40_IA, were purchased from JPT Peptide Technologies (Germany). Two recombinant Aβ variants were purchased as lyophilized powder from AlexoTech AB (Umeå, Sweden), namely the Aβ_40_ peptide and the Aβ_40_(H6A, H13A, H14A) triple-mutant, which in the following is referred to as the Aβ_40_(NoHis) mutant. The Aβ_40_ peptide was purchased also as uniformly single-labeled with ^15^N isotopes. All peptides were stored at −80 °C until use. Before measurements, they were dissolved in 10 mM NaOH, and then sonicated in an ice bath for 5 minutes to avoid pre-formed aggregates. The samples were then diluted in either sodium phosphate buffer at pH 7.3, or in MES buffer at pH 7.3 or 5.1. The peptide concentrations were determined from the weight of the dry powder.

### 3.2 Preparation of Aβ_42_ oligomers

Treatment of Aβ_42_ peptides with low concentrations (≤ 7 mM) of SDS, i.e. below the critical micelle concentration for SDS which is 8.2 mM in water at 25 °C ^111^, leads to formation of stable and homogeneous Aβ_42_ oligomers of certain sizes and conformations ^112-114^. To prepare such oligomers, size exclusion chromatography (SEC) was initially used to purify synthetic Aβ_42_ peptides into monomeric form. First, 1 mg of lyophilized Aβ_42_ powder was dissolved in 250 mL of DMSO. Next, a Sephadex G-250 HiTrap desalting column (GE Healthcare, Uppsala) was equilibrated with 5 mM NaOH solution (pH=12.3), and washed with a solution of 10–15 mL of 5 mM NaOD, pD=12.7 ^115^. The peptide solution in DMSO was applied to the column, followed by injection of 1.25 mL of 5 mM NaOD. Collection of peptide fractions in 5 mM NaOD on ice was started at a 1 mg/ mL flow rate. Ten fractions of 1 mL volumes were collected in 1.5 mL Eppendorf tubes. The absorbance for each fraction at 280 nm was measured with a NanoDrop instrument (Eppendorf, Germany), and peptide concentrations were determined using a molar extinction coefficient of 1280 M^-1^cm^-1^ for the single Tyr in Aβ_42_ ^116^. The peptide fractions were flash-frozen in liquid nitrogen, covered with argon gas on top in 1.5 mL Eppendorf tubes, and stored at –80 °C until used. SDS-stabilized Aβ_42_ oligomers of two well-defined sizes (approximately tetramers and dodecamers) were prepared according to a previously published protocol ^112^, but in D_2_O, at a 4-fold lower peptide concentration, and without the original dilution step ^114^. The reaction mixtures (100-120 μM Aβ_42_ in PBS, containing either 0.05% (1.7 mM) SDS or 0.2% (6.9 mM) SDS) were incubated together with 0–1000 µM uranyl acetate at 37 °C for 24 hours, and then flash-frozen in liquid nitrogen and stored at –20 °C for later analysis.

### 3.3 Thioflavin T (ThT) aggregation Kinetics

To monitor the effect of uranyl ions on Aβ_40_ aggregation kinetics, a FLUOstar Omega microplate reader (BMG LABTECH, Germany) was used. Samples containing 20 µM Aβ_40_ wildtype, 20 mM MES buffer pH 7.3, 50 µM Thioflavin T, and different concentration of uranyl acetate (i.e., 0 µM, 0.04 µM, 0.2 µM, 0.4 µM, 2 µM, and 20 µM) were added to a 384-well plate, with 35 µl of sample in each well. Thioflavin T is a benzothiazole dye that increases in fluorescence upon binding to amyloid aggregates ^117^ and is therefore used to monitor formation of amyloid aggregates. The ThT dye was excited at 440 nm, and ThT fluorescence emission at 480 nm was measured every five minutes. Before each measurement, the plate was shaken for 140 seconds at 200 rpm. The samples were incubated for a total of 15 hours, and the assay was repeated three times with four replicates of each condition. To determine kinetic aggregation parameters, the data was fitted to equation 1:

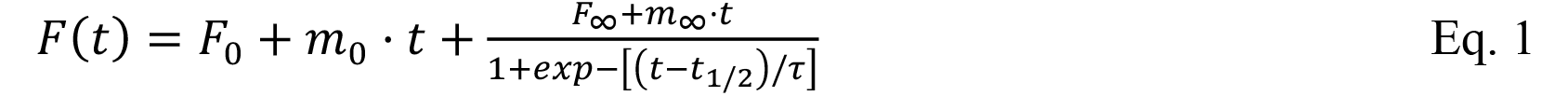

Here, F_0_ and F_∞_ are the intercepts of the initial and final fluorescence intensity baselines, m_0_ and m_∞_ are the slopes of the initial and final baselines, t_½_ is the time needed to reach halfway through the elongation phase (i.e., aggregation half-time), and τ is the elongation time constant ^117^.

### 3.4 Transmission electron microscopy (TEM) imaging

Negative staining TEM images were recorded for Aβ_40_ peptides that had aggregated under the same conditions as in the ThT fluorescence studies (above). Thus, 20 µM of Aβ_40_ in 20 mM MES buffer, pH 7.3, was incubated for 20 hours on a thermo shaker at 37 °C and 300 rpm, together with 0 µM, 0.2 µM, 2 µM, and 20 µM of uranyl acetate. Then, samples of 5 µL were put on copper grids of 200 µm mesh size, which were covered with Pioloform film upon which a carbon layer had been deposited and then glow-discharged with a Leica EM ACE600 carbon coater (Leica Microsystems, Germany). The Aβ_40_ samples were absorbed to the grids for 5 mins, rinsed with Milli-Q water two times, and then stained for 2 mins with 2% aqueous solution of uranyl acetate. Next, the excess stain was removed with filter paper, and the samples were left to air-dry. A digital Orius SC1000 camera was used to record TEM images in a FEI Tecnai G2 Spirit electron microscope (FEI, The Netherlands) operating at 120 kV accelerating voltage.

### 3.5 Circular dichroism (CD) spectroscopy measurements of secondary structure

Circular dichroism (CD) was carried out using a Chirascan CD spectrometer from Applied Photophysics Ltd. (U.K.). Sample containing 600 µl of 10 µM Aβ_40_ peptide in 20 mM phosphate buffer, at either pH 7.3 or pH 5.1, were placed in a quartz cuvette with 2 mm pathlength. CD spectra were recorded at 20 °C between 192 nm and 250 nm, using steps of 0.5 nm and a sampling time of 5 s per data point. Then, small volumes of uranyl acetate were titrated to the samples in steps of 2 µM, 6 µM, 16 µM, 56 µM, and finally 256 µM. The total increase of volume upon addition of the uranyl acetate was less than 3 %. All data was processed with the Chirascan Pro-Data v.4.4.1 software (Applied Photophysics Ltd., U.K.), including smoothing with a ten points smoothing filter.

50 mM SDS detergent was added to some of the samples, as SDS micelles constitute a simple model for bio-membranes ^118-119^. Aβ peptides are known to insert their central and C-terminal segments as α-helices into SDS micelles, while the N-terminal Aβ segment remains unstructured outside the micelle surface ^17^, ^118^. Because the critical micelle concentration for SDS is 8.2 mM in water at 25 °C ^111^, micelles clearly formed under the experimental conditions. With approximately 62-65 SDS molecules per micelle ^120^, 50 mM SDS yields a micelle concentration slightly below 1 mM, i.e. much higher than the concentration of Aβ peptides. This means that each micelle will generally contain no more than one Aβ peptide, which effectively prevents Aβ aggregation and ensures that uranyl interactions are with monomeric Aβ peptides.

### 3.6 Fluorescence measurements of dityrosine formation

Fluorescence emission spectra between 330 and 480 nm (excitation at 315 nm) were recorded at room temperature with a Jobin Yvon Horiba Fluorolog 3 fluorescence spectrometer (Longjumeau, France), for two samples of 10 µM Aβ_40_ peptide in 20 mM MES buffer, pH 7.3. One sample contained 100 µM uranyl acetate, to investigate possible effects of uranyl ions on dityrosine formation. The control sample contained 100 µM of the chelator EDTA, to remove any free metal ions. All measurements were conducted in triplicate, using a quartz cuvette with 4 mm path length and containing 0.7 mL liquid sample. Spectra were recorded after 0 and 24 hrs of incubation, during which the samples were kept at room temperature without agitation or other treatment.

### 3.7 Nuclear magnetic resonance (NMR) spectroscopy

Nuclear magnetic resonance (NMR) spectroscopy experiments were conducted on a Bruker Avance spectrometer operating at 500 MHz and being equipped with a cryoprobe for increased sensitivity. Uranyl acetate was titrated to 92 μM monomeric ^15^N-labelled Aβ_40_ peptides in 20 mM MES buffer (90/10 H_2_O/D_2_O) at either pH 7.3 or pH 5.1 at 5 °C. NMR spectra (2D ^1^H,^15^N-HSQC) were recorded during the titrations and then processed and evaluated in the Topspin software (v. 3.2), using already published assignments for HSQC cross-peaks of Aβ_40_ in buffer at neutral pH ^121-123^ or at acidic pH ^124^.

### 3.8 Binding affinity measurements via tyrosine fluorescence quenching

The binding affinities between uranyl ions and Aβ_40_ peptides were evaluated via the quenching effect of uranyl on the intrinsic fluorescence of Tyr10, the only natural fluorophore in wild-type Aβ peptide. Fluorescence measurements were conducted on a Jobin Yvon Horiba Fluorolog 3 fluorescence spectrophotometer (Longjumeau, France), using a quartz cuvette with 4 mm path length. The fluorescence emission intensity of samples containing 20 µM Aβ peptides in 20 mM MES buffer, at either pH 7.3 or pH 5.5 and without or with 50 mM SDS detergent present, were measured at 305 nm (excitation wavelength 276 nm) at 20 °C. Aliquots of uranyl acetate (stock concentrations of 1, 2 and 10 mM) were titrated to the samples, and the Tyr10 fluorescence intensity was plotted against the concentration of UO_2_^2+^ ions. Apparent dissociation constants (K_D_^App^) were determined by fitting the plots to equation 2:

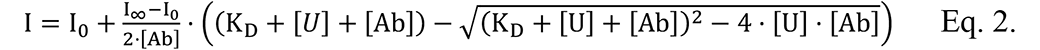

where I_0_ is the initial fluorescence intensity with no added UO_2_^2+^ ions, I_∞_ is the steady-state intensity at the end of the titration, [Ab] is the protein concentration, and [U] is the concentration of added UO_2_^2+^ ions ^125-126^.

### 3.9 Blue native polyacrylamide gel electrophoresis

Homogeneous solutions of oligomers of 80-100 μM Aβ_42_ peptides ^114^ prepared (as described in section 3.2) in the presence of different concentrations of uranyl acetate (0–1000 µM) were analyzed with blue native polyacrylamide gel electrophoresis (BN-PAGE) using the Invitrogen system (ThermoFisher Scientific, USA). Thus, 4–16% Bis-Tris Novex gels (ThermoFisher Scientific, USA) were loaded with 10 μL of Aβ_42_ oligomer samples alongside the Amersham High Molecular Weight calibration kit for native electrophoresis (GE Healthcare, USA). The gels were run at 4°C using the electrophoresis system according to the Invitrogen instructions (ThermoFisher Scientific, USA), and then stained with the Pierce Silver Staining kit according to the manufacturer’s instructions (ThermoFisher Scientific, USA). BN-PAGE was chosen for analysis instead of SDS-PAGE to avoid disruption of the SDS-stabilized and non-cross linked Aβ_42_ oligomers by the high (>1%) SDS concentrations used in SDS-PAGE sample buffers ^127^.

### 3.10 Infrared spectroscopy

Fourier-transformed infrared (FTIR) spectra of the SDS-stabilized Aβ_42_ oligomers (prepared as described in section 3.2) were recorded in transmission mode on a Tensor 37 FTIR spectrometer (Bruker Optics, Germany) equipped with a sample shutter and a liquid nitrogen-cooled MCT detector. The unit was continuously purged with dry air during the measurements. 8–10 μL of the 80 μM Aβ_42_ oligomer samples, prepared (as described in section 3.2) with different concentrations of uranyl acetate (0–1000 µM), were put between two flat CaF_2_ discs separated by a 50 μm plastic spacer covered with vacuum grease at the periphery. The assembled discs were mounted into the sample position of a sample shuttle inside the instrument’s sample chamber, while a metal grid (used as the background) was positioned in the reference holder. The sample shuttle was used to measure sample and reference spectra without opening the chamber. The samples were allowed to sit for at least 15 minutes after closing the chamber lid, to avoid interference from water vapor. FTIR spectra were recorded at room temperature in the 1900–800 cm^−1^ range, with 300 scans for both background and sample spectra, using a 6 mm aperture and at a resolution of 2 cm^−1^. The light intensities above 2200 cm^−1^ and below 1500 cm^−1^ were blocked with respectively a germanium filter and a cellulose membrane (Baldassarre & Barth, 2014). The spectra were analyzed and plotted with the OPUS 5.5 software, and second derivatives were calculated with a 17 cm^−1^ smoothing range.

## 4. Results

### 4.1 ThT fluorescence measurements of Aβ_40_ aggregation kinetics

The fluorescence intensity of ThT, a common marker for amyloid material ^117^, was monitored when samples of 20 µM Aβ_40_ peptides in 20 mM MES buffer, pH 7.3, were incubated under quiescent conditions for 15 hrs, at 37 °C, together with different concentrations (0 µM; 0.04 µM; 0.2 µM; 0.4 µM; 2 µM; and 20 µM) of uranyl acetate (Fig. 1).

Fitting Eq. 1 to the ThT fluorescence curves in Fig. 1 produced the kinetic parameters T_lag_ and t_½_ (Table 1). For the three highest concentrations of uranyl acetate, i.e. 0.4 µM, 2 µM, and 20 µM, the ThT kinetics curves are mostly flat, and fitting Eq. 1 to these curves was not possible. The flat shape of the three curves indicate that no or very little amyloid material has formed, although for the 0.4 µM uranyl sample, small amounts of ThT-binding material has begun to form after approximately nine hours. It is obvious that the amount of amyloid material formed at the end of the measurements, i.e., ΔThT, strictly decreases with the amount of added uranyl acetate (Fig. 1; Table 1). The two samples with 0 µM and 0.04 µM uranyl both have aggregation half-times around three hours, and lag times around two hours, under the experimental conditions (Table 1). The aggregation of the 0.2 µM uranyl sample is clearly slower, with a half-time of almost seven hours, and a lag time around five hours. Given the visually estimated lag time of around nine hours for the 0.4 µM uranyl sample, and that the 2 µM and 20 µM uranyl samples show no obvious signs of aggregation after 15 hrs, uranyl ions clearly increase the lag time for Aβ_40_ aggregation.

**Figure 1.**
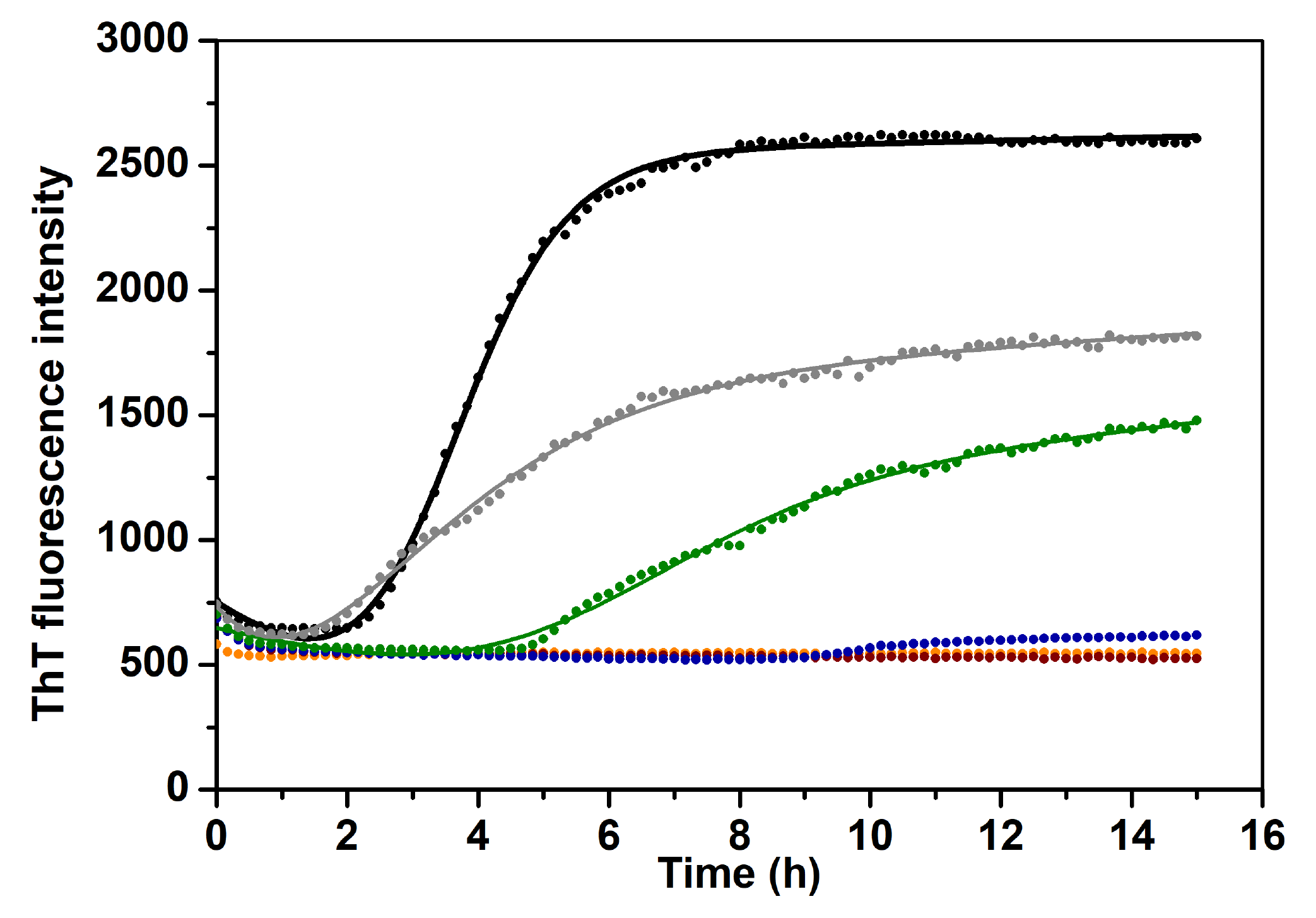
Monitoring the fibril formation of Aβ_40_ peptides via Thioflavin T fluorescence. 20 µM Aβ_40_ was incubated at 37 °C in 20 mM MES buffer, pH 7.3, together with different concentrations of uranyl acetate: 0 µM (black); 0.04 µM (grey); 0.2 µM (green); 0.4 µM (blue); 2 µM (red); and 20 µM (orange).

**Table 1.**
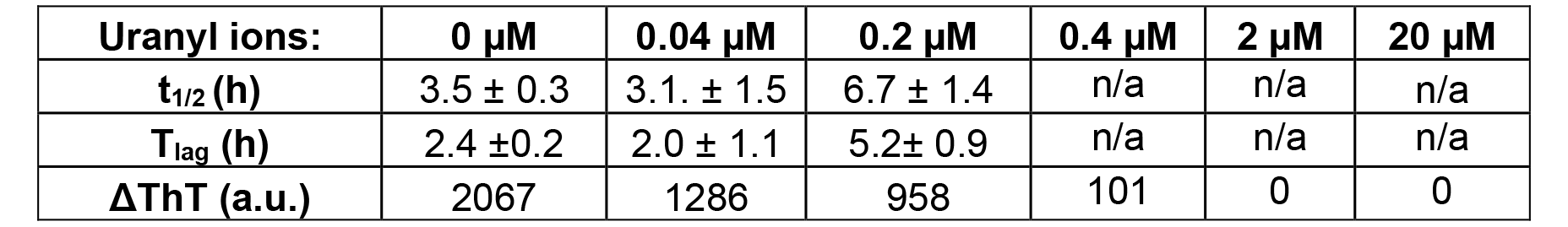
Kinetic parameters for Aβ_40_ aggregation in the presence of different concentrations of uranyl acetate, derived from the ThT fluorescence curves shown in Fig. 1. Aggregation halftime (t_1/2_) and lag time (T_lag_) were obtained from fitting the ThT curves to Eq. 1. The increase in ThT fluorescence (ΔThT) is presented in arbitrary fluorescence units (a.u.).

### 4.2 TEM images of aggregated Aβ_40_ peptides

Negative staining TEM images were recorded for 20 μM Aβ_40_ peptide, incubated for 20 hours with different concentrations of uranyl acetate, i.e., 0 µM, 0.2 µM, 2 µM, and 20 µM (Fig. 2). The uranyl additions correspond to uranyl:Aβ_40_ ratios of 1:100, 1:10, and 1:1. The control sample without UO_2_^2+^ ions displays numerous amyloid fibrils that are several microns long, together with occasional larger clumps and smaller aggregates which may be proto-fibrils (Fig. 2A). This is in line with previous in vitro studies of Aβ_40_ aggregates ^8^, ^23^, ^117^. The Aβ_40_ samples incubated together with 0.2 µM of uranyl acetate display amyloid fibrils of similar size and shape (Fig. 2B). In the presence of 2 µM uranyl acetate there are fewer fibrils and they are shorter, i.e., only a few microns long (Fig. 2C). At the highest concentration of uranyl acetate, i.e. 20 µM, no fibrils have formed at all (Fig. 2D). Instead, the Aβ_40_ peptides have aggregated into large amorphous clumps, which co-exist with smaller particles (Fig. 2D). These results show that UO_2_^2+^ ions inhibit formation of Aβ amyloid fibrils in a concentration-dependent manner, with complete inhibition at stoichiometric uranyl:Aβ_40_ ratios.

**Figure 2.**
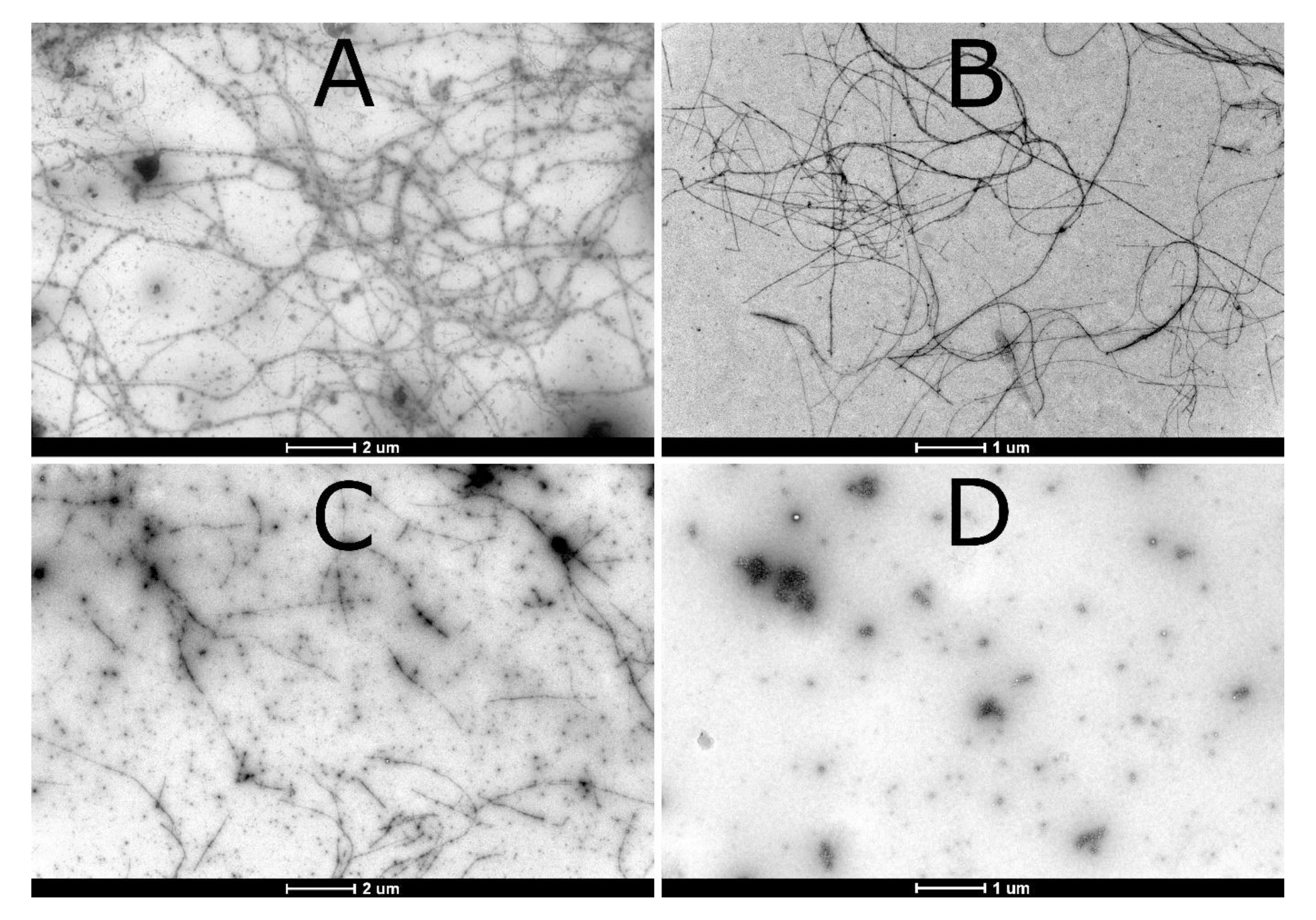
TEM images for aggregates of 20 µM Aβ_40_ in 20 mM MES buffer, pH 7.3, incubated for 20 hours on a thermo shaker at 37 °C and 300 rpm, together with different concentrations of uranyl acetate: A) 0 µM; B) 0.2 µM; C) 2 µM; and D) 20 µM. The white scale bars are either 2 µm (A and C) or 1 µm (B and D).

### 4.3 NMR spectroscopy on uranyl binding to Aβ_40_ monomers

High-resolution NMR experiments were conducted to investigate if there were residue-specific molecular interactions between uranyl ions and monomeric Aβ_40_ peptides. Fig. 3 shows 2D ^1^H,^15^N-HSQC spectra for the amide cross-peak region for 92 µM ^15^N-labeled Aβ_40_ peptides, at either pH 7.3 or pH 5.1, recorded before and after addition of uranyl acetate.

**Figure 3.**
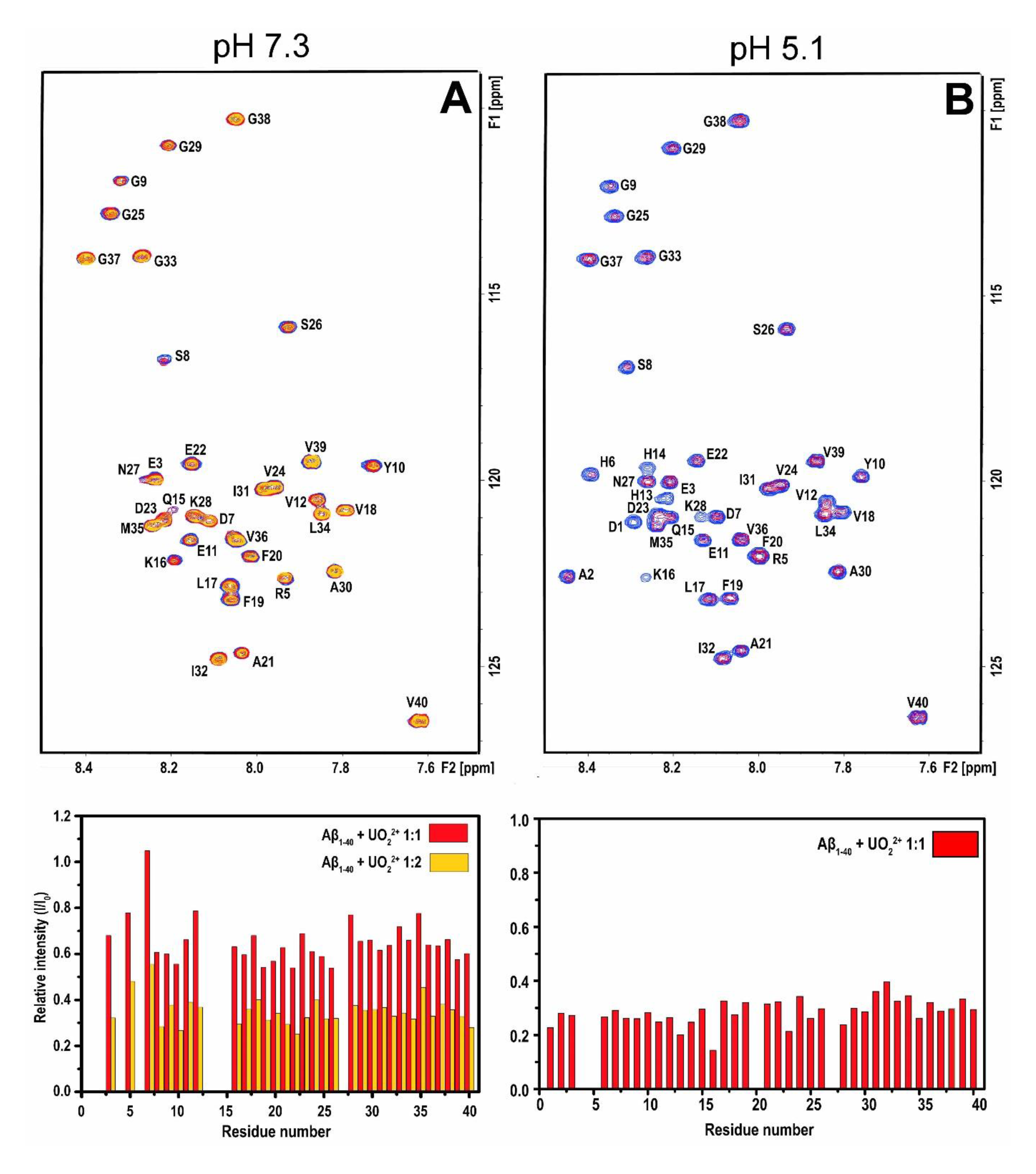
2D NMR ^1^H,^15^N-HSQC-spectra at 5 °C showing titrations of uranyl acetate to 92 µM monomeric ^15^N-labeled Aβ_40_ peptides in 20 mM MES buffer at either pH 7.3 (A) or pH 5.1 (B). Blue crosspeaks: 84 µM Aβ_40_ only. Red crosspeaks: 1:2 uranyl:Aβ_40_. Yellow crosspeaks (pH 7.3 only): 1:1 uranyl:Aβ_40_. The peak intensities in the bar charts show ratios between the crosspeak intensities with added uranyl ions relative to the intensities before addition of uranyl ions, i.e. I/I_0_.

At pH 7.3, addition of first 46 µM and then 92 µM uranyl ions (1:2 and 1:1 uranyl:Aβ_40_ ratio, respectively) induces a concentration-dependent loss of amide cross-peak intensity (Fig. 3A). The intensity loss is uniformly distributed across the peptide sequence, which shows that the uranyl ions do not bind to specific residues of the Aβ_40_ monomer. Instead, the observed binding is likely driven by general electrostatic interactions between the cationic uranyl ions and the anionic Aβ peptides. The loss of cross-peak intensity is probably caused by several factors, such as paramagnetic quenching effects ^94-95^ and intermediate (on the NMR time-scale) chemical exchange between free Aβ_40_ peptides and the Aβ_40_·uranyl complex, similar to the effects observed when Cu(II), Ni(II), and Zn(II) ions bind to Aβ peptides ^124^, ^128-130^. In addition, the uranyl ions likely promote formation of Aβ aggregates that either precipitate out of the solution or are too large and heterogeneous to produce distinct NMR signals ^124^, ^130^. As no NMR signals are observed for the Aβ_40_ aggregates, nothing can be concluded about the binding configurations for uranyl ions in complex with the aggregates.

At pH 5.1, addition of uranyl ions again induces a uniform loss of amide cross-peak intensity (Fig. 3B). Unexpectedly, the effect is stronger at pH 5.1 than at pH 7.3: addition of 46 µM uranyl acetate (1:2 uranyl:Aβ_40_ ratio) at pH 5.1 decreases the cross-peak intensity approximately as much as addition of 92 µM uranyl acetate (1:1 ratio) at pH 7.3 (Fig. 3). This suggests stronger binding of the uranyl ions at acidic pH. Compared to the spectrum at neutral pH (Fig. 3A), NMR crosspeaks for a few additional residues (e.g., D1, A2, H6, H13, and H14) became visible at acidic pH (Fig. 3B), probably due to slower proton exchange at this pH ^124^.

### 4.4 Fluorescence measurements of uranyl·Aβ binding affinity

Uranyl ions were found to quench the intrinsic fluorescence of the Tyr10 residue in the Aβ peptide, similar to e.g. Cu(II) ions ^125^, ^131-132^. Measurements of the Tyr10 fluorescence during titrations with uranyl acetate were therefore used to quantify binding affinities for Aβ·UO_2_^2+^ complexes under different conditions. The resulting titration/binding curves are shown in Fig. 4. Fitting Eq.2 to these curves produced the apparent dissociation constants (K_D_^App^) shown in Table 2.

**Figure 4.**
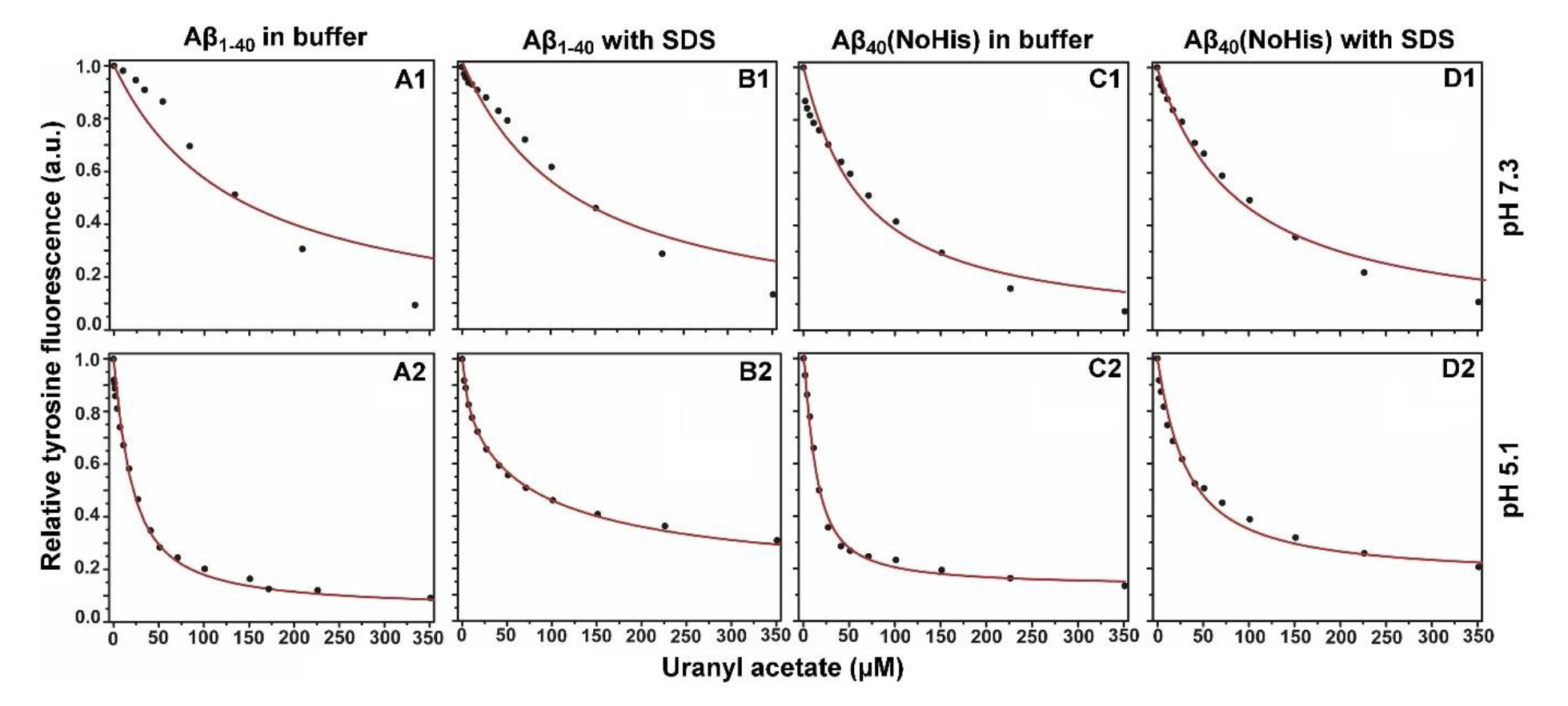
Fluorescence intensity curves for Aβ residue Tyr10, showing titrations of 20 µM Aβ_40_ peptide (A and B) or 20 µM Aβ_40_(NoHis) mutant (C and D) with uranyl acetate in 20 mM MES buffer, either at pH 7.3 (1st row) or at pH 5.1 (2nd row). For B and D, micelles of 50 mM SDS were added to the samples.

**Table 2.** Apparent dissociation constants (K_D_^App^) in µM for uranyl·Aβ complexes, obtained by fitting fluorescence titration curves to Eq.2 (Fig. 4; three replicates per condition).

| | A$\beta_{40}$ | | | | A$\beta_{40}(\text{NoHis})$ | | | |
| --- | --- | --- | --- | --- | --- | --- | --- | --- |
|  | 1 | 2 | 3 | Average | 1 | 2 | 3 | Average |
| pH 7.3 | 130 | 141 | 148 | $140 \pm 8$ | 62 | 86 | 79 | $76 \pm 11 \mu\text{M}$ |
| pH 7.3 + 50mM SDS | 206 | 252 | 251 | $236 \pm 22$ | 119 | 124 | 116 | $120 \pm 4 \mu\text{M}$ |
| pH 5.1 | 14.0 | 13.7 | 21.3 | $16.3 \pm 4$ | 2.3 | 2.8 | 4.0 | $3.0 \pm 1 \mu\text{M}$ |
| pH 5.1 + 50mM SDS | 21.8 | 21.7 | 27.4 | $23.6 \pm 3$ | 22.3 | 30.7 | 27.9 | $27.0 \pm 4 \mu\text{M}$ |

For Aβ_40_ at pH 7.3, the titration data clearly deviate from the binding model, both for the measurements in buffer only (Fig. 4:A1) and in the presence of SDS micelles (Fig. 4:B1). Because of these deviations, the derived dissociation constants should not be trusted. As Eq.2 is based on a model that assumes a single binding site, a possible explanation for this deviation is binding of uranyl ions to multiple locations on the Aβ_40_ peptide, under these conditions. At pH 5.1, the binding curves follow the model very well (Fig. 4:A2 and B2), yielding reliable K_D_^App^ values of 16.3 µM for Aβ_40_ in buffer, and 23.6 µM for Aβ_40_ bound to SDS micelles (Table 2). The main difference from the measurements at neutral pH is that the three His residues in the Aβ peptide become protonated at lower pH, as their pKa values are around 6.8 ^133^. Because protonated His residues are unlikely to interact with positive ions such as UO_2_^2+^, it is possible that binding interactions between uranyl ions and uncharged His residues are responsible for the deviations from single-site binding scheme observed at pH 7.3 (Fig. 4:A1 and B1).

The binding curves for the Aβ_40_(NoHis) mutant at pH 7.3 follow the model rather well (Fig. 4:C1 and D1), producing K_D_^App^ values of 76 µM in buffer, and 120 µM in the presence of SDS micelles (Table 2). At pH 5.1 the overlap with the fitted curve is even better (Fig. 4:C2 and D2), and here the K_D_^App^ values are 3.0 µM in buffer, and 27 µM in the presence of SDS micelles. With these results, some comparisons can be made.

For both peptide versions, the uranyl ions display stronger binding at low pH. Even though reliable binding constants could not be obtained for Aβ_40_ at pH 7.3, it is clear from the measurement data that binding is overall stronger at pH 5.1 (Fig. 4:A and B). Addition of SDS micelles generally makes the binding weaker (Table 2). The strongest binding is observed for the Aβ_40_(NoHis) mutant at pH 5.1 (3.0 µM), while the weakest binding is observed for Aβ_40_ at pH 7.3. It therefore appears that neutral (i.e., non-protonated) His residues might interfere with uranyl binding, under the experimental conditions used.

### 4.5 CD spectroscopy on Aβ secondary structure

CD spectroscopy was used to investigate possible effects of uranyl ions on the secondary structure of Aβ_40_ peptides, both in aqueous buffer and in the presence of SDS micelles that mimic a membrane environment. The CD spectra for Aβ_40_ monomers in aqueous buffer display spectra with minima around 196-198 nm (Fig. 5:B and E), which is typical for a random coil conformation. At pH 5.1, addition of uranyl ions to Aβ_40_ induces a two-step structural transition, as the CD signal changes in a concentration-dependent manner around an isodichroic point at approximately 214 nm (Fig. 3E). The difference spectrum (Fig. 5F) shows that the structural transition is a conversion from random coil to β-sheet. At pH 7.3, the structural effect induced by the uranyl ions is weaker, and it is unclear if an isodichroic point is present (Fig. 5B). The difference spectrum is not conclusive, but might represent β-sheet conformation (Fig. 5C).

**Figure 5.**
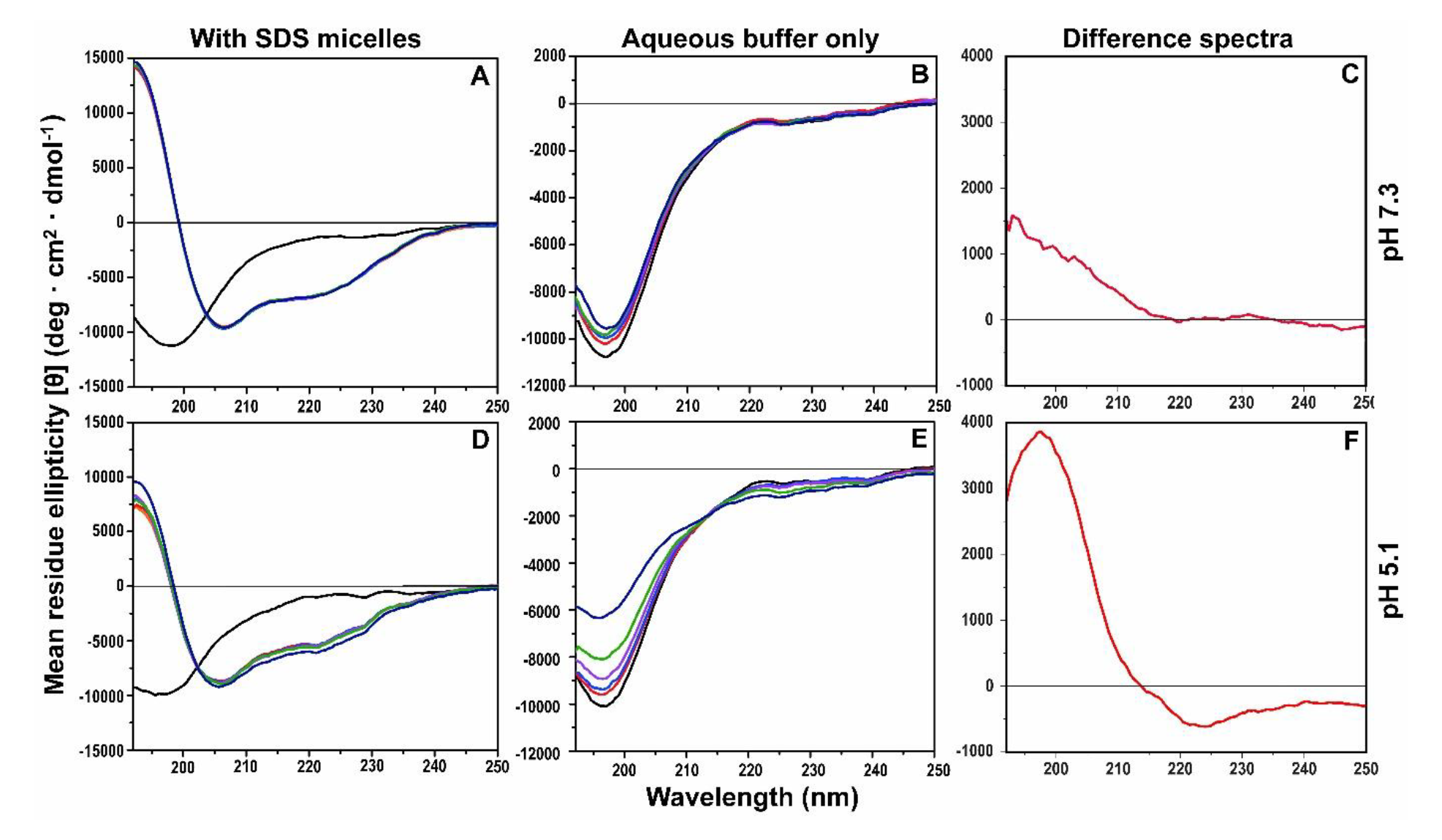
CD spectroscopic titrations of 10 µM Aβ_40_ with uranyl acetate in 20 mM sodium phosphate buffer at 20 °C, carried out in the presence of 50 mM SDS at pH 7.3 (A) or pH 5.1 (D), or in aqueous buffer only at pH 7.3 (B) or pH 5.1 (E). First, spectra were recorded for Aβ_40_ in buffer, for all samples (black lines). Then, 50 mM SDS detergent was added to the samples in A and D (orange lines). To all samples, uranyl acetate was added in steps of 2 µM, 6 µM, 16 µM, 56 µM, and 256 µM (ruby red, ultramarine blue, plum purple, lime green, and navy blue lines, respectively). For the titrations in aqueous buffer, shown in (B) and (E), the difference spectra shown in (C) and (F) were created by subtracting the measurements with 256 µM added uranyl from the measurements without uranyl, at pH 7.3 (C) and at pH 5.1 (F).

In the presence of SDS micelles, the CD signals for Aβ_40_ peptides display minima around 208 and 222 nm (Fig. 5:A and D), which is characteristic for α-helical secondary structure. This is in line with previous reports that the central and C-terminal Aβ regions adopt α-helical conformations in membrane-like environments ^17^, ^118-119^, ^131^. At pH 7.3, addition of uranyl ions induces no changes at all in the Aβ_40_ CD spectrum (Fig. 5A). At pH 5.1, however, the uranyl ions induce systematic changes in the CD spectra around an isodichroic point at approximately 203 nm (Fig. 5D). This is the same wavelength as where the CD spectrum for Aβ_40_ peptides in aqueous solution (i.e., random coil structure) crosses the CD spectrum for Aβ_40_ peptides in SDS micelles (i.e., α-helical structure), when no uranyl ions are present (Fig. 5D). Taken together, these observations indicate that uranyl ions at pH 5.1 induce a structural conversion from α-helix to random coil in Aβ_40_ peptides in SDS micelles. Also the changes in CD intensity at other wavelengths are consistent with formation of random coil structure, even though the overall changes are small and difficult to interpret conclusively (Fig. 5D). The overall larger structural effects induced by uranyl ions at acidic pH, compared to neutral pH, both in aqueous solution (Fig. 5:B and E) and in SDS micelles (Fig. 5:A and D), support the NMR (Fig. 3) and fluorescence (Fig. 4) spectroscopy results that the uranyl ions bind stronger to Aβ_40_ peptides at acidic pH.

### 4.6 Dityrosine cross-link formation

Fluorescence measurements were carried out to investigate if uranyl ions could induce formation of covalent dityrosine crosslinks in Aβ peptides, via generation of reactive oxygen species (ROS), similar to what has been observed for redox-active Cu(II) ions ^32^, ^134-136^ and Ni(II) ions ^129^. Two samples of Aβ_40_ peptides were studied. The control sample contained 100 µM EDTA, to remove possible contaminating metal ions that could promote dityrosine formation. The other sample contained 100 µM uranyl acetate. For both samples, the fluorescence spectra are virtually identical before and after 24 hrs of incubation (Fig. 6). Specifically, no peak around 410 nm, indicative of dityrosine ^136-137^, has emerged. The signal around 350 nm is a water Raman peak, which is useful for reference purposes, as it does not change with the sample composition. These results clearly show that uranyl ions do not induce formation of dityrosine crosslinks, under the experimental conditions employed here.

**Figure 6.**
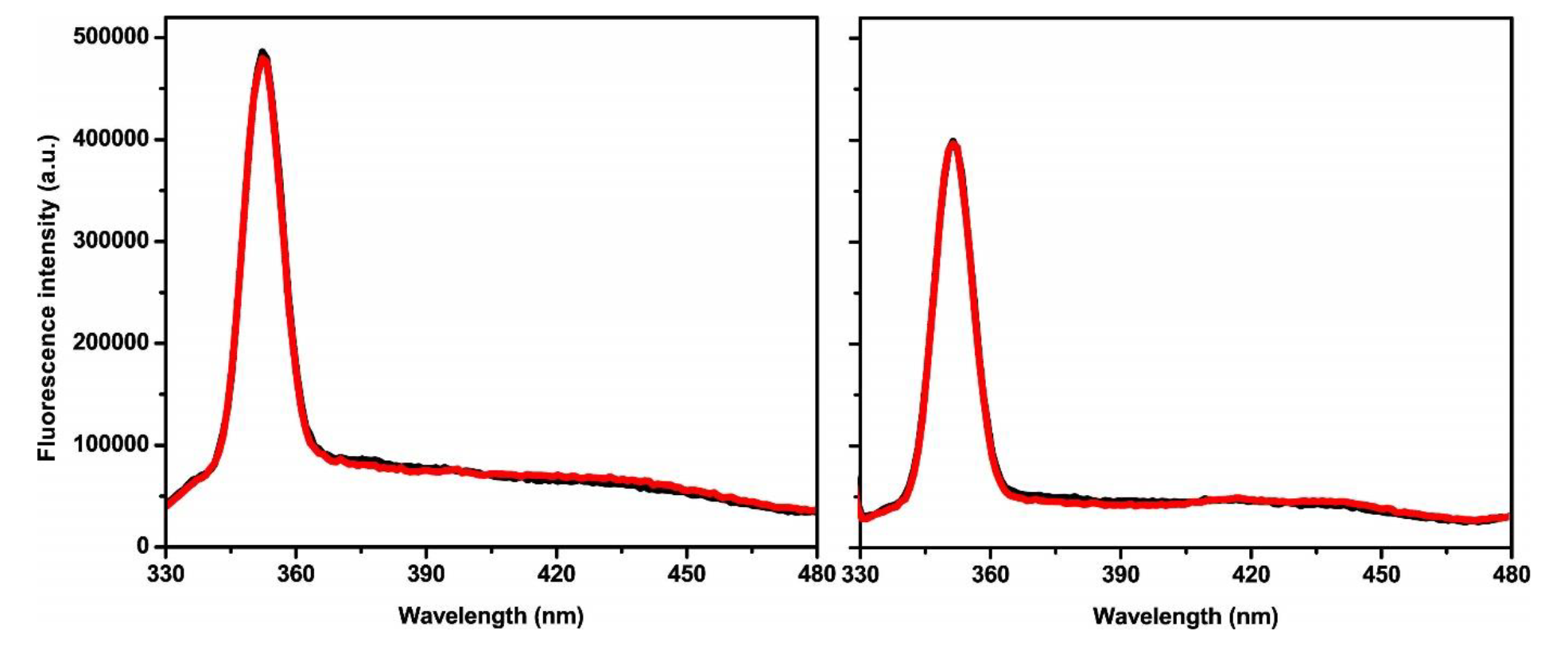
Fluorescence emission spectra of 10 µM Aβ_40_ peptides in 20 mM MES buffer, pH 7.3, incubated with 100 µM EDTA (A) or with 100 µM uranyl acetate (B). Black line – 0 hrs; red line – 24 hrs.

### 4.7 BN-PAGE analysis of Aβ_42_ oligomer formation and stability

BN-PAGE analysis and FTIR analysis (below) were used to investigate possible effects of uranyl ions on Aβ oligomer formation. Because Aβ_40_ peptides do not form stable oligomers, Aβ_42_ peptides together with SDS detergent were used to create oligomers that remained stable for over a week. The Aβ_42_ oligomers were created with two different concentrations of stabilizing SDS molecules, 80-100 µM of Aβ_42_ peptides, and 0-1000 µM uranyl acetate as described in section 3.2.

The BN-PAGE analysis shows that in 0.05% SDS (1.7 mM) without uranyl acetate, larger oligomers with a molecular weight (MW) around 55-60 kD are formed (Fig. 7, Lane 3). These larger oligomers, abbreviated AβO_0.05%SDS_, most likely contain twelve Aβ_42_ peptides and display a globular morphology, which is why they sometimes are called globulomers ^112^. In the presence of different concentrations of uranyl acetate, no well-defined uniform oligomers were observed. Instead, the bands on the BN-PAGE gel appear smeared, indicating a heterogeneous size distribution of the formed oligomers (Fig. 7, Lanes 4-6). This disruptive effect is very strong already at the lowest concentration (10 µM) of added uranyl ions.

**Figure 7.**
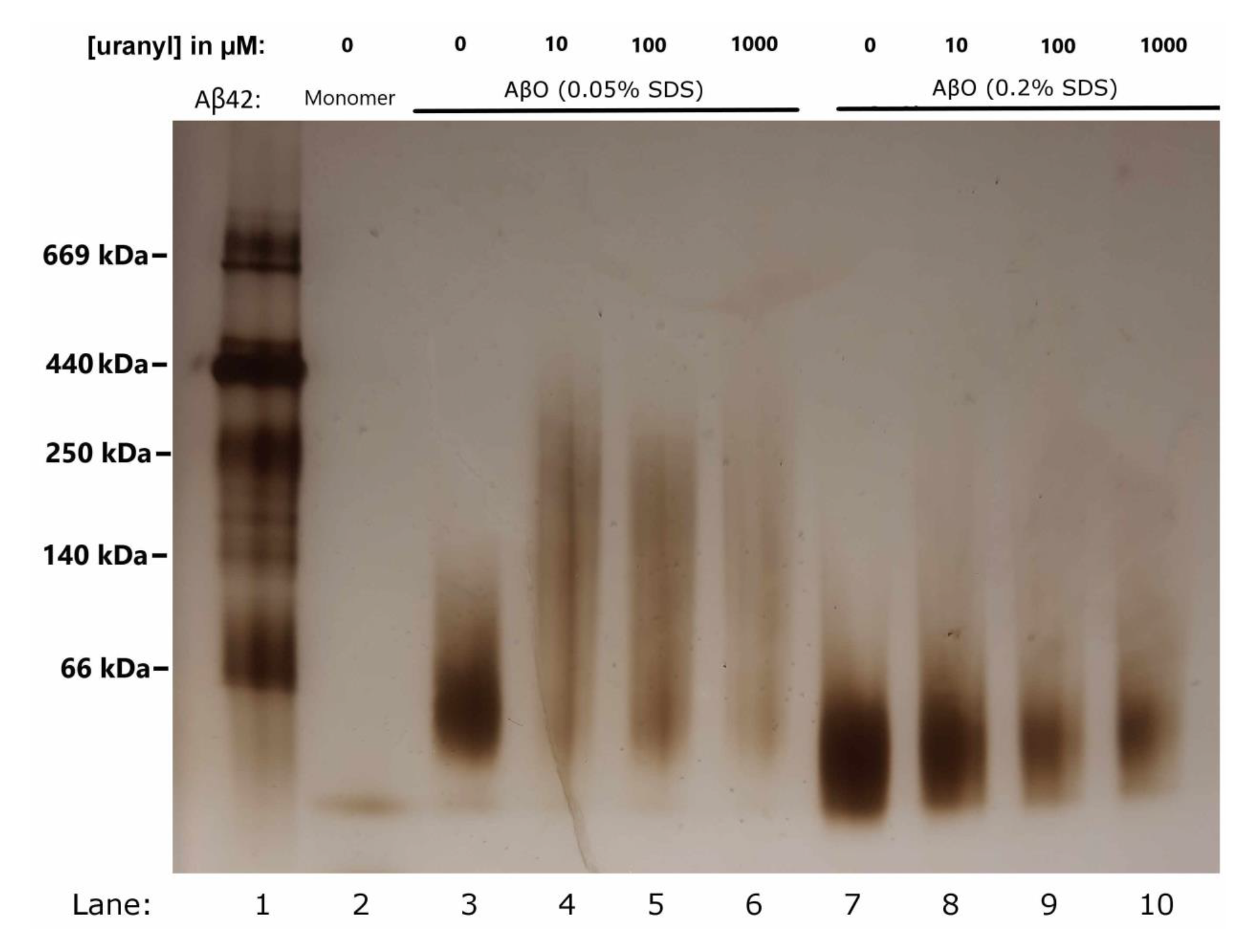
BN-PAGE gel showing the effects of different concentrations of uranyl acetate on the formation of SDS-stabilized Aβ_42_ oligomers (formed with 80-100 µM peptide). Lane 1: Reference ladder. Lane 2: Aβ_42_ monomers. Lanes 3-6: AβO_0.05%SDS_ oligomers (likely dodecamers) prepared with 0, 10, 100, and 1000 µM uranyl ions, respectively. Lanes 7-10: AβO_0.2%SDS_ oligomers (likely tetramers) prepared with 0, 10, 100, and 1000 µM uranyl ions, respectively.

In 0.2% SDS (6.9 mM), small Aβ_42_ oligomers with a MW around 16-20 kDa are formed (Fig. 7, Lane 7). These oligomers, abbreviated AβO_0.2%SDS_, likely contain a large fraction of tetramers ^112^, ^114^. Preparation of these oligomers in the presence of increasing uranyl concentrations (10, 100, and 1000 µM) yields oligomer bands that are increasingly weaker, indicating gradually lower amounts of stable AβO_0.2%SDS_ oligomers (Fig. 7, Lanes 8-10). Thus, for formation of the smaller AβO_0.2%SDS_ oligomers, the effect of uranyl ions is less disruptive and clearly concentration-dependent, compared to the effect on the larger AβO_0.05%SDS_ oligomers (Fig. 7).

### 4.8 FTIR spectroscopy reflecting Aβ_42_ oligomer structure

The secondary structures of Aβ_42_ oligomers formed with different SDS and uranyl concentrations were studied with FTIR spectroscopy, where the amide I region (1700- 1600 cm^-1^) is very sensitive to changes in the protein backbone conformation, including differences in β-sheet structures ^138-140^. When prepared in absence of uranyl acetate, both types of Aβ_42_ oligomers (i.e., AβO_0.05%SDS_ and AβO_0.2%SDS_) produce two major bands in the amide I region: a high-intensity band at 1630-1629 cm^-1^ and a smaller one at about 1685 cm^-1^ ^114^. These IR bands are typical for anti-parallel β-sheet structures ^141-143^.

For AβO_0.2%SDS_ oligomers in PBS buffer, the β-sheet main band is observed at 1630.2 cm^-1^ (Fig. 8). When these oligomers are formed in the presence of 0.01 mM, 0.1 mM, and 1 mM of uranyl acetate, the β-sheet band position shifts to 1629.5, 1629.8, and 1629.4 cm^-1^, respectively (Fig. 8:A and B). These shifts generally suggest formation of larger oligomers with more extended β-sheet structures.

**Figure 8.**
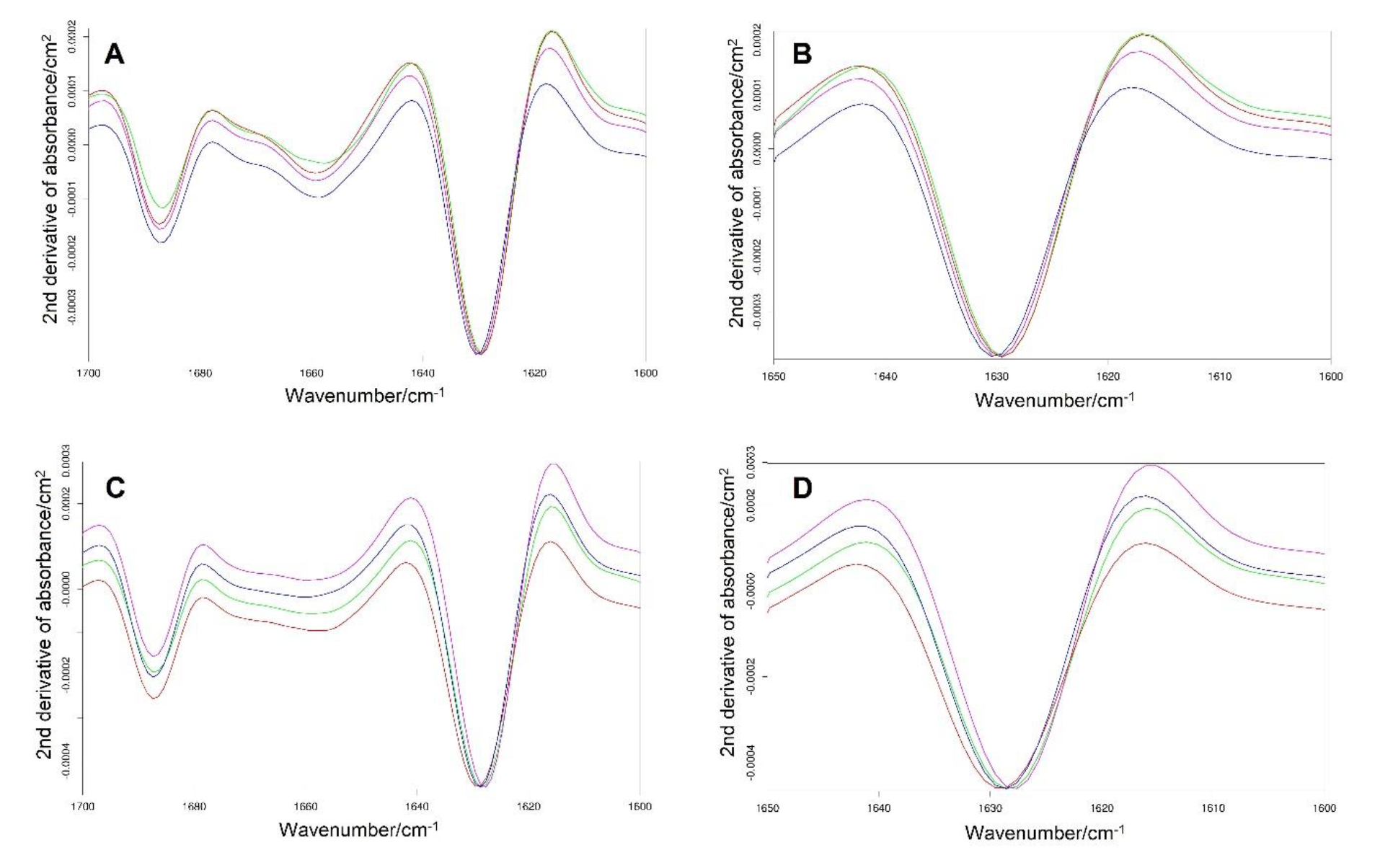
Second derivatives of infrared absorbance spectra for 80-100 µM SDS- stabilized Aβ_42_ oligomers formed in absence (blue) and presence of 0.01 mM (red), 0.1 mM (purple), and 1 mM (green) uranyl acetate. The results are shown for smaller Aβ_42_ oligomers formed with 0.2% SDS (A and B) and for larger Aβ_42_ globulomers formed with 0.05% SDS (C and D).

For AβO_0.05%SDS_ oligomers, the β-sheet main band is observed at 1628.7 cm^-1^ without uranyl acetate, but downshifts to around 1628.3 cm^-1^ in presence of 1 mM uranyl acetate (Fig. 8:C and D). Thus, the uranyl-induced downshift is slightly larger than for the AβO_0.2%SDS_ oligomers, indicating that the AβO_0.05%SDS_ oligomer conformation is more sensitive to uranyl-induced effects.

## 5. Discussion

Uranium is a well-known neurotoxicant ^34^, but the underlying molecular mechanisms and a possible role of U in neurodegenerative diseases remain unclear ^44^, ^86^. For Alzheimer’s disease, several studies have investigated how Aβ peptides interact with various metal ions ^28^, ^42^, ^129^, ^144-147^ and with small cationic molecules ^26^, ^148^. We here interpret the current results on Aβ-uranyl interactions in the light of this earlier work.

### 5.1. Effects of uranyl ions on Aβ_40_ aggregation

The ThT fluorescence curves (Fig. 1) and TEM images (Fig. 2) show that uranyl ions have a concentration-dependent inhibitory effect on the fibrillization of Aβ_40_ peptides at physiological pH. Instead of forming proper amyloid fibrils, the Aβ_40_ peptides form non-fibrillar amorphous aggregates (clumps) already at a 1:1 uranyl/Aβ_40_ ratio (Fig. 2). Thus, the uranyl ions do not prevent the Aβ_40_ peptides from aggregating, but rather direct the aggregation process towards non-fibrillar end-products. Similar effects on the Aβ aggregation pathways have previously been observed for other small cationic molecules and heavy metal ions such as Hg(II), Ni(II), Pb(II) and Pb(IV) ^42^, ^129^, ^148^. As small oligomeric aggregates of Aβ peptides are considered to be the main toxic species in AD pathology ^2^, ^11^, the finding that uranyl ions can modulate Aβ aggregation might be relevant for understanding AD progression and pathogenesis.

### 5.2 Binding of uranyl ions to Aβ_40_ peptides

The NMR results show that uranyl ions have no obvious residue-specific binding to Aβ_40_ peptides (Fig. 3). Because of the strong covalent O=U=O bonds in the uranyl ion, it is mainly the central U(VI) atom that interacts with other molecules ^149^. Thus, the uranyl-Aβ binding is probably mediated via non-specific electrostatic interactions between the positive U(VI) atom and negative Aβ residues, such as Asp1, Asp7, Asp23, Glu3, Glu11, and Glu22. This is similar to Aβ binding to Mn(II) and Pb(II) ions ^3^, ^144^, but different from numerous small cationic species that display residue-specific interactions with Aβ peptides, such as polyamines ^26^ and Cu(II), Ni(II), Pb(IV) and Zn(II) ions ^3^, ^124^, ^128-129^.

The results of the NMR measurements indicate that the Aβ binding of uranyl ions is stronger at acidic pH (Fig. 3), which is confirmed by the fluorescence measurements of uranyl-Aβ binding affinity (Fig. 4), and also by the CD spectroscopy results (Fig. 5). At pH 5.1, the apparent binding affinity of uranyl ions is 16.3 ± 4 µM to Aβ_40_ peptides, and 3.0 ± 1 µM to the Aβ_40_(NoHis) mutant (Table 2). At neutral pH the binding is about a magnitude weaker (Table 2), and also less well-defined (Fig. 4). This is counter-intuitive. The main chemical difference at lower pH is that the Aβ His residues become protonated, as they have pKa values around 6.8 in short peptides ^133^. It stands to reason that Aβ peptides with positively charged His residues should be less prone to interact with positive metal ions, as has been shown for e.g. Cu(II) and Zn(II) ions ^124^.

The Tyr10 fluorescence measurements (Fig. 4) show that the uranyl·Aβ binding affinity increases at lower pH values, and also when the three His residues are replaced by Ala residues (Table 2). In fact, the weakest binding is observed for Aβ_40_ at pH 7.3, and the strongest binding is observed for the Aβ_40_(NoHis) mutant at pH 5.1 (Table 2). This trend is observed for Aβ samples both in aqueous buffer and in SDS micelles. It strongly suggests that the presence of uncharged His sidechains interferes with uranyl binding to Aβ peptides (Table 2). This tentative conclusion is supported by the shape of the fluorescence data curves, which are more uniform and better fit the single-binding-site model (Eq. 2), when the uncharged His residues H6, H13, and H14 are replaced with Ala residues in the Aβ_40_(NoHis) mutant (Fig. 4). In addition, at acidic pH the uranyl binding is stronger both to the wt Aβ_40_ peptide and to the Aβ_40_(NoHis) mutant. In the latter case, the effect is clearly not caused by His protonation, as these residues are missing in the mutant peptide. Thus, the His residues are not the only factor responsible for the unusual binding properties of the uranyl·Aβ complex. The molecular details of this complex should therefore be further explored in future studies.

For all conditions and Aβ variants, the binding affinities decrease slightly when SDS micelles are present in the sample (Fig. 4, Table 2). This is similar to the effect of SDS micelles on Aβ binding to other metal ions such as Cu(II), Hg(II), and Ni(II) ^42^, ^129^, ^131^. Aβ peptides are known to insert only their central and C-terminal regions into an SDS micelle, while the negatively charged N-terminal region is positioned outside the micelle where it is free to interact with e.g. cations ^129^, ^131^. Thus, it appears that Aβ peptides can (sometimes) bind uranyl ions mainly via their N-terminal part.

### 5.3 Effects of uranyl ions on Aβ_40_ peptide structure

In aqueous buffer at pH 5.1, the CD spectroscopy measurements show that uranyl ions induce a two-state structural transition in the Aβ_40_ peptides, from random coil to β- sheet structure (Fig. 5:E and F). At pH 7.3, a weaker structural transition is observed, that might be a similar conversion into β-sheet structure (Fig. 5:B and C). Earlier studies have shown that such structural changes can be induced also by metal ions such as Cu(II), Ni(II), and Zn(II) ^124^, ^129^. Because Aβ aggregates typically consist of peptides in β-sheet conformation ^16^, this type of β-sheet structure formation likely promotes Aβ aggregation.

In the presence of SDS micelles, which constitute a simple membrane model ^118-119^ where the central and C-terminal Aβ regions insert themselves into the micelle and adopt α-helical conformations, ^17^, ^119^, uranyl ions have no effect on the Aβ_40_ peptide conformation at pH 7.3 (Fig. 5A). However, a weak structural transition is observed at pH 5.1, possibly involving formation of random coils (Fig. 5D). Previous studies have found that metal ions that bind to Aβ peptides mainly via the His residues, such as Cu(II), Ni(II), and Zn(II) ions, can induce altered coil-coil interactions in Aβ peptides positioned in SDS micelles ^129^, ^131^. No such alterations in the α-helical structure were observed upon addition of uranyl ions (Fig. 5:A and D).

No dityrosine cross-links were observed after Aβ_40_ peptides had been incubated together with uranyl acetate (Fig. 4). Earlier studies have shown that redox-active metal ions such as Cu(II) and Ni(II) can induce formation of dityrosine crosslinks in amyloid peptides, via redox-cycling of e.g. the Cu(I)/Cu(II) redox pair, which generates harmful oxygen radicals via Fenton-like chemistry ^32^, ^129^, ^136^. For short peptides with only one Tyr residue in the amino acid sequence, such as Aβ and amylin, dityrosine formation must involve two peptides, which then become linked to form a dimer ^136^. As Cu(II) and Ni(II) bind Aβ peptides mainly via the His6, His13, and His14 residues, it is likely that these metal ions can promote Aβ aggregation by coordinating multiple His residues from more than one Aβ peptide ^146^, ^150^. Cu(II) and Ni(II) ions may therefore promote dityrosine formation not only by creating oxygen radicals, but also by connecting two Aβ molecules and positioning their Tyr10 residues close to each other. It is known that the U(VI) ion in uranyl can be reduced to lower valency states such as U(IV) under appropriate reducing conditions ^149^, ^151^. Thus, as uranyl acetate has been found to induce oxidative stress in isolated cells ^92^, the uranyl ions are likely redox-active under physiological conditions. We therefore speculate that the reason why uranyl ions do not promote dityrosine formation in Aβ_40_ peptides (Fig. 4) is the weak and non-specific binding under the experimental conditions used (Figs. 7 and 8; Table 2).

### 5.4 Effects of uranyl ions on Aβ_42_ oligomers

Although most of the measurements carried out in this study were performed on Aβ_40_ monomers, it is of interest to investigate also the possible effects of uranyl ions on Aβ oligomers. Because Aβ_40_ peptides do not form stable oligomers, BN-PAGE and FTIR studies were carried out on Aβ_42_ oligomers stabilized by SDS detergent. The BN-PAGE experiments clearly show that uranyl ions interfere with formation of homogeneous Aβ_42_ oligomers (Fig. 7). This is further supported by the FTIR measurements, where the position of the β-sheet main band is downshifted when increasing concentrations of uranyl ions are present during oligomer formation (Fig. 6). The uranyl effect is more pronounced on the larger and mainly dodecameric Aβ_42_ oligomers, which are formed in presence of 0.05% SDS. The spectral changes observed with uranyl ions are consistent with the previously observed effects of Ni(II) ions ^129^. However, only 10 μM of divalent uranyl ions (Fig. 5) but 500 mM of divalent Ni(II) ions ^129^ are required for full disruption of homogenous AβO_0.05%SDS_ oligomers, even though Ni(II) ions display stronger binding affinity for Aβ peptides than uranyl ions, at neutral pH (Figs. 7 and 8; Table 2). In general, the effects of uranyl ions on Aβ oligomerization more resemble those of other transition metal ions, including Ni ^129^, rather than those of monovalent alkali ions, such as Li ^145^. Such conclusions are consistent with the theoretical findings regarding the relative propensities of different metal ions for interactions with polypeptides ^152^.

## 6. Conclusions

Uranyl ions, UO_2_^2+^, bind to Aβ_40_ peptides *via* non-specific electrostatic interactions, with an apparent binding affinity of 16.3 ± 4 µM at pH 5.1. Uranyl binding is weaker and less uniform at neutral pH, possibly because of interference from His sidechains. The uranyl ions inhibit Aβ fibrillization and oligomer formation in a concentration-dependent manner, and induce structural changes in both Aβ monomers and oligomers. A general toxic mechanism of uranyl ions could be to modulate protein folding, misfolding, and aggregation.

## Acknowledgments

This work was supported by grants from the Estonian Research Council to MP (no. PRG1506) and to PP (no. 1289), from the Magnus Bergvall foundation to SW, from the Kamprad Research Foundation, the Ulla-Carin Lindquist Foundation for ALS Research and the Karolinska Institutet IMM strategic grants to PMR, from the Swedish Research Council and the Brain Foundation in Sweden to AG, and from the National Science Centre in Poland to MK (no. 2017/27/B/ST4/00485).

## Conflicts of interest

AG, JJ, and SW are shareholders in CellPept Sweden AB. Neither this company, nor the funding organizations, had any role in the design of the study; in the collection, analyses, or interpretation of data; in the writing of the manuscript; or in the decision to publish the results.

## Notes

### Competing Interest Statement

The authors have declared no competing interest.

